# Controlled pH Alteration Enables Guanine Accumulation and Drives Crystallization within Iridosomes

**DOI:** 10.1101/2024.07.20.604036

**Authors:** Zohar Eyal, Anna Gorelick-Ashkenazi, Rachael Deis, Yuval Barzilay, Yonatan Broder, Asher Perry Kellum, Neta Varsano, Michal Hartstein, Andrea Sorrentino, Ifat Kaplan-Ashiri, Katya Rechav, Rebecca Metzler, Lothar Houben, Leeor Kronik, Peter Rez, Dvir Gur

**Affiliations:** Department of Molecular Genetics, Weizmann Institute of Science; Rehovot, 7610001, Israel; Department of Biomolecular Sciences, Weizmann Institute of Science; Rehovot, 7610001, Israel; Department of Chemical Research Support, Weizmann Institute of Science; Rehovot, 7610001, Israel; Department of Materials and Interfaces, Weizmann Institute of Science; Rehovot, 7610001, Israel; Mistral Beamline, ALBA Synchrotron Light Source, Cerdanyola del Vallès, 08290, Spain; Department of Physics and Astronomy, Colgate University, New York, 13346, USA; Department of Physics, Arizona State University, Arizona, 85287, USA

## Abstract

Many animals exhibit remarkable colors produced by the constructive interference of light reflected from arrays of intracellular guanine crystals. These systems are utilized for various purposes, including vision, camouflage, communication, and thermal regulation. Each guanine crystal forms within a membrane-bound organelle called an iridosome, where precise control over crystal formation occurs. While the presence of guanine crystals in iridosomes is well-documented, the mechanisms facilitating the accumulation of water-insoluble guanine and driving its crystallization remain unclear. Here, we employ advanced imaging and spectroscopy techniques to characterize the maturation of iridosomes in zebrafish iridophores during development. Using cryo-electron microscopy, we found that amorphous guanine accumulates in early-stage iridosomes. Synchrotron-based soft X-ray microscopy studies revealed that, unlike mature crystals, the accumulated guanine is initially in its protonated state. Live imaging with a pH sensor demonstrated that early-stage iridosomes are acidic and that their pH gradually approaches neutrality during maturation. Additionally, the application of a V-ATPase inhibitor reduced the acidity of iridosomes and significantly decreased crystal formation, suggesting the involvement of V-ATPase in regulating the organelle pH. Our findings reveal new insights into the molecular mechanisms facilitating guanine accumulation and crystallization within iridosomes, emphasizing the pivotal role of pH alternations in the precise formation of biogenic crystals.

## Introduction

Iridophores are specialized pigment cells that encapsulate reflective guanine crystals^1,2^, playing a critical role in the vibrant and dynamic coloration observed in a variety of organisms, including fish, reptiles, amphibians, cephalopods, and certain spiders^3-11^. These cells enhance color variation and facilitate adaptive color changes for camouflage, signalling, and vision^6,10,12-14^. Biogenic guanine crystals are formed within specialized membrane-bound organelles within iridophores, known as iridosomes^15^. The process leading to crystal formation, predominantly taking place during the early developmental stages of iridophores, involves the synthesis and trafficking of guanine precursors to the iridosome, their concentration within the organelle, and the eventual crystallization into bio-organic structures with highly controlled properties^6,8,16,17^.

Several factors influence the guanine crystallization process, including supersaturation, temperature, solvent properties, the presence of impurities, agitation, and importantly, pH. pH significantly impacts the ionization and solubility of guanine, which is insoluble under neutral conditions and only dissolves in extremely acidic or alkaline environments^18,19^. Manipulating the pH of crystallization solutions *in vitro* has been shown to control guanine’s crystal phases, polymorphism, and size, reflecting the stability of different guanine tautomers under varying pH conditions^18^.

Iridophores, along with melanophores—which also originate from the neural crest^1,20^-contain organelles that were suggested to be lysosome-related organelles (LROs) family^16,21-23^. These organelles are distinguished by their specific cargoes and functions^22-24^. While melanosomes have been well-studied for their development from endosomal structures and their role in melanin synthesis^22,25-27^, the detailed biochemical dynamics and structural development of iridosomes remain less explored.

While the exact mechanism of crystal formation differs among organisms, biogenic guanine crystals have been shown to form via non-classical mechanisms^28-31^. In this process, amorphous guanine initially accumulates within the iridosome until templated crystallization occurs on preassembled scaffolds^31,32^. However, a key question remains: What are the mechanisms facilitating the accumulation of water-insoluble guanine and driving its crystallization?

Here we utilize zebrafish (*Danio rerio*) iridophores as a model system to investigate the process taking place in the iridosomes before crystal formation. By employing advanced imaging and spectroscopic techniques, we aim to delineate the mechanisms through which the microenvironment within iridosomes affects guanine accumulation and crystallization. We show that the organelle pH plays a crucial role in this process. Initially, the early-stage iridosome is highly acidic, promoting the accumulation of protonated guanine and the assembly of molecular scaffolds. As the organelle matures, the neutralization of pH drives the nucleation and subsequent crystal growth.

## Results

To investigate the early stages of guanine accumulation and the key factors that lead to its crystallization, we focused our investigation on the iridophores of the zebrafish, as a model system. Zebrafish employ guanine crystals for various functions. These crystals, distributed throughout their skin and eyes, are crucial in shaping skin color and patterns, enabling camouflage, functioning as light barriers, and improving visual acuity, particularly in dim environments (**Fig. 1A-B**)^14,33-38^. Notably, the initial appearance of iridophore crystals in zebrafish larvae occurs around 44 hours post-fertilization (hpf)^39^. This timing makes the zebrafish an excellent model for studying guanine accumulation and subsequent crystal formation.

**Figure 1:**
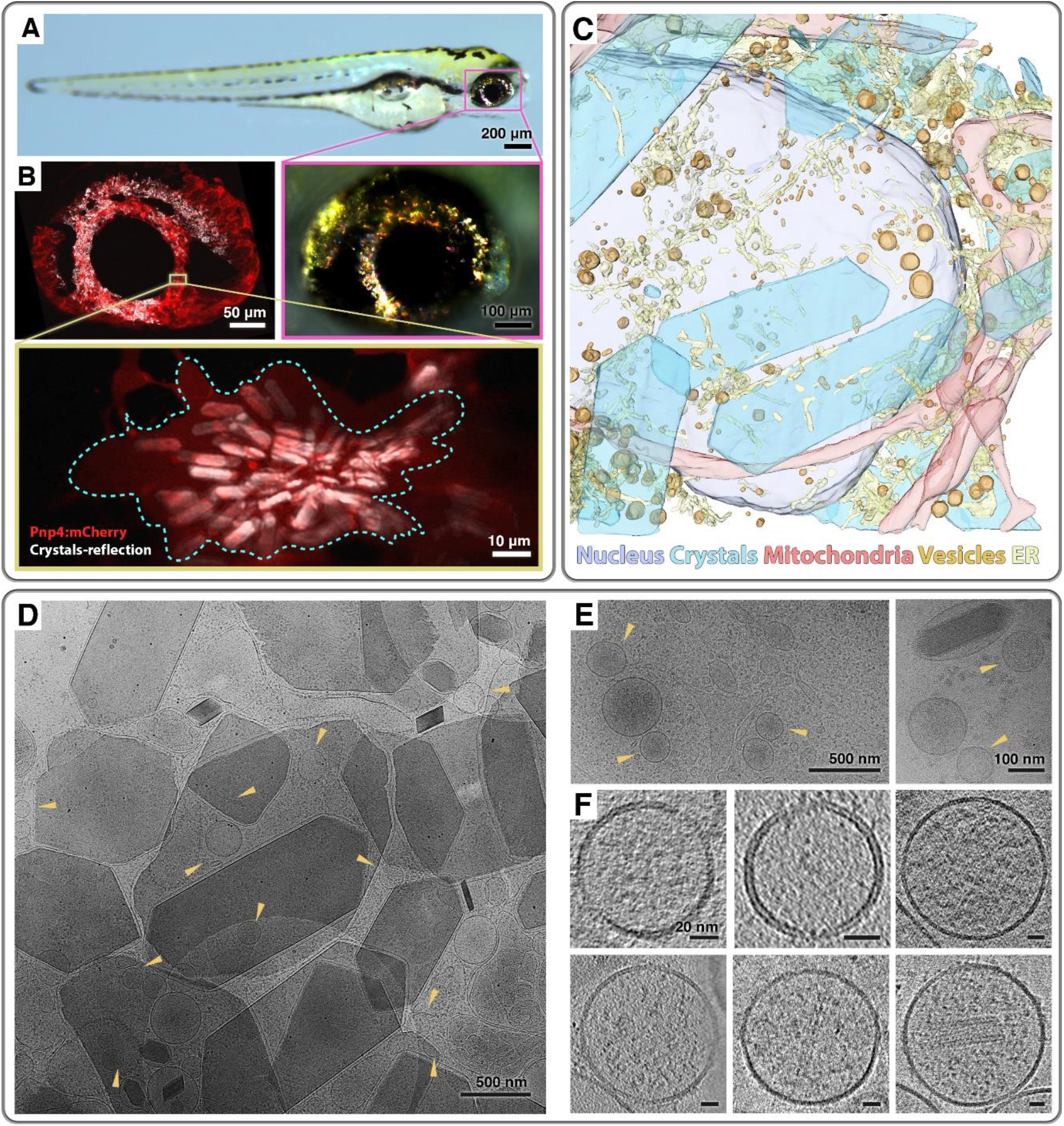
Intracellular electron-dense vesicles are abundant in zebrafish larva iridophores. **(A)** Incident light images of a zebrafish larva at 5 dpf containing guanine crystals in its eyes and skin. Inset shows higher magnifications of the eye (pink). **(B)** Confocal images of zebrafish larva eye at 72 hpf showing labeled iridophores (Tg(*pnp4a:palmmCherry*), red) and crystal reflection (white); high magnification of a single iridophore (yellow frame). **(C)** A 3D surface representation of Cryo-FIB-SEM data of eye iridophore from 72 hpf larva, where hundreds of nanometer size vesicles are segmented in yellow, and crystals are segmented in blue. **(D, E)** Cryo-TEM micrographs of isolated iridophores from zebrafish larva, yellow arrowheads mark a variety of hundreds of nanometer scale intracellular vesicles with high-density content. **(F)** Cryo-ET reconstruction images of different vesicle populations found in iridophores cells.

To capture the initial stages that take place prior to crystal formation within their natural context, we examined eye iridophores in zebrafish larvae *in situ* using cryogenic focused ion beam scanning electron microscopy (cryo FIB-SEM). This approach revealed numerous vesicles, each measuring hundreds of nanometers in diameter, located adjacent to iridosomes at various stages of development (**Fig. 1C and Movie S1**). To enhance the visualization of these vesicles and their contents, we employed cryogenic transmission electron microscopy (cryo-TEM). This was achieved by cryo-fixing iridophores, isolated by fluorescence-activated cell sorting (FACS), through rapid plunging in liquid ethane, which allowed for higher-resolution imaging in close to native conditions. Through this technique, we observed vesicles ranging from 100 to 500 nanometers in diameter. Notably, some of these vesicles exhibited different levels of contrast, suggesting the presence of electron-dense material akin to that found in the crystals (**Fig. 1D-E**). In several cases, vesicles containing dense material were also found to contain fibers (**Fig. 1F**). These fibers have been previously proposed to act as scaffolds, facilitating the templated nucleation of crystals^31,32^.

To identify the composition within the electron-dense intracellular vesicles, we employed a combination of cryo-scanning electron microscopy and energy dispersive X-ray spectroscopy (cryo-SEM-EDS). This method was applied to larvae that underwent high-pressure freezing and freeze-fracturing for *in situ* imaging and spectroscopy (**Fig. 2A)**. Analysis revealed that the iridosomes, which encapsulate guanine crystals, were notably rich in nitrogen (**Fig. 2B-C and Fig. S1A-G)**. This aligns with the chemical structure of guanine, where nitrogen constitutes five of its sixteen atoms **(Fig. S2A)**. Intriguingly, we found that intracellular vesicles, similar in size to those detected earlier via cryo-TEM, also exhibited significant nitrogen enrichment (**Fig. 2B-C and Fig. S1F-G**).

**Figure 2:**
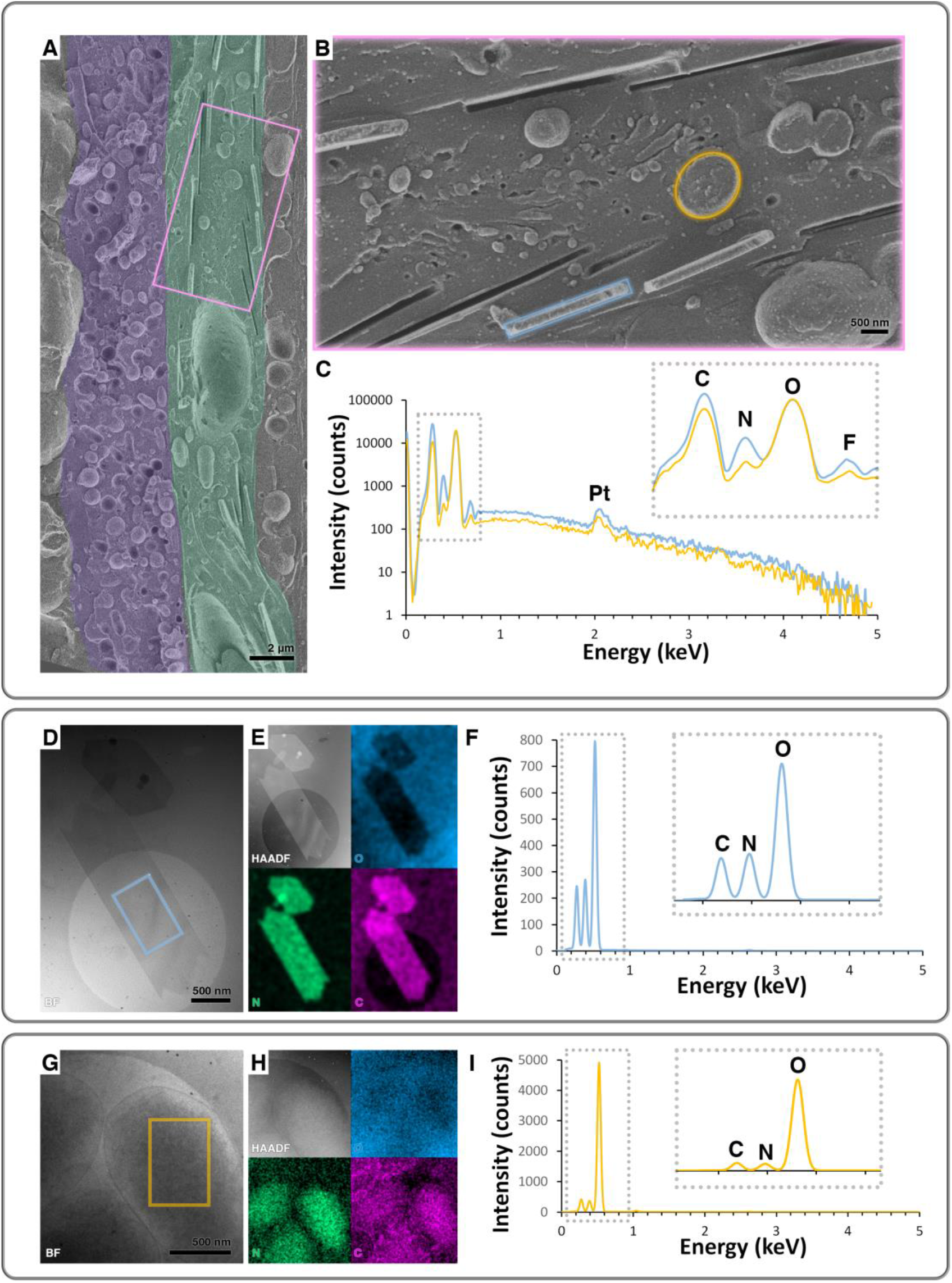
Cryo-EDS imaging reveals a high concentration of nitrogen within crystals and vesicles. **(A)** Cryo-SEM image of freeze-fractured surface of iridophores (green) and melanophores (purple) within the zebrafish larva eye at 72 hpf. **(B)** A higher magnification view of an iridophore from A (pink rectangular). **(C)** EDS spectrum taken from the marked regions in B: intracellular vesicle (orange circle) and crystal (cyan rectangular). Inset shows a closer view of the 0-1 keV region. Both spectra show the presence of fluorine (F), platinum (Pt), oxygen (O), nitrogen (N) and carbon (C) in these areas. **(D, G)** Cryo-STEM bright field image of guanine crystals (D) and vesicles (G) from isolated iridophores of 72 hpf larvae. **(E, H)** Cryo-STEM dark field and EDS maps of oxygen (blue), nitrogen (green) and carbon (magenta). **(F, I)** EDS corresponds to the marked regions of interest in D and G: intracellular vesicle (orange frame) and crystal (cyan frame). Insets show a closer view of the 0-1 keV region.

To further substantiate these findings and to achieve better spatial and spectral resolution, we next utilized cryo-scanning transmission electron microscopy and energy dispersive X-ray spectroscopy (cryo-STEM-EDS). This technique confirmed our previous observations: both the guanine crystals (**Fig. 2D-F**) and the high-contrast vesicles (**Fig. 2G-I**) were enriched in nitrogen. To correlate the morphological information with crystallographic features, we investigated the high-contrast vesicles using cryogenic 4D scanning transmission electron microscopy (cryo-4D-STEM) to collect an electron diffraction pattern from every point the beam raster traverses (**Fig. S1F-G**). We found that the crystals exhibited a diffraction pattern that correlated to the *bc* plane of anhydrous β-guanine^40^ (**Fig. S1H**). In contrast, the high-contrast vesicles showed no crystalline spots in a diffraction pattern typical for amorphous compounds, despite the high sensitivity of the method for nanometer-sized crystals (**Fig. S1I**). These findings suggest that the vesicles, like the crystals, also contain guanine or other related nitrogen-rich substrate. However, unlike the crystals, the material within the vesicles does not exhibit the organized structure characteristic of the crystals.

To further elucidate the composition of the nitrogen-rich material in the intracellular vesicles, and to further investigate the connection between nitrogen accumulation and guanine crystallization, we employed a combination of synchrotron-based cryo soft X-ray tomography (cryo-SXT) and X-ray absorption near edge structure (XANES). The correlative use of these advanced techniques facilitates the visualization of entire isolated cells while concurrently allowing for chemical characterization with high spatial and spectral resolution using XANES-microscopy^41,42^ (**Fig. 3A**) of plunge-frozen isolated iridophores at various stages of maturation (**Fig. 3**). This was achieved by performing tilt series within the ‘water window’ energy range. Within this energy range, carbon and nitrogen exhibit high absorption levels, while oxygen and, consequently, water is almost transparent. This results in carbon and nitrogen-rich substances appearing darker in transmission images, whereas the water-abundant cytosol appears lighter (**Movie S2**). Moreover, X-ray attenuation properties allow the acquisition of data from samples up to 15 µm thick, permitting the imaging of whole cells that have been disaggregated from larvae. The reconstructed tomograms reveal various cellular compartments with enhanced contrast compared to the cytoplasm, including the cell membrane, nucleus, guanine crystals, and diverse vesicles potentially containing nitrogen-rich material (**Fig. 3 and Movie S3**).

**Figure 3:**
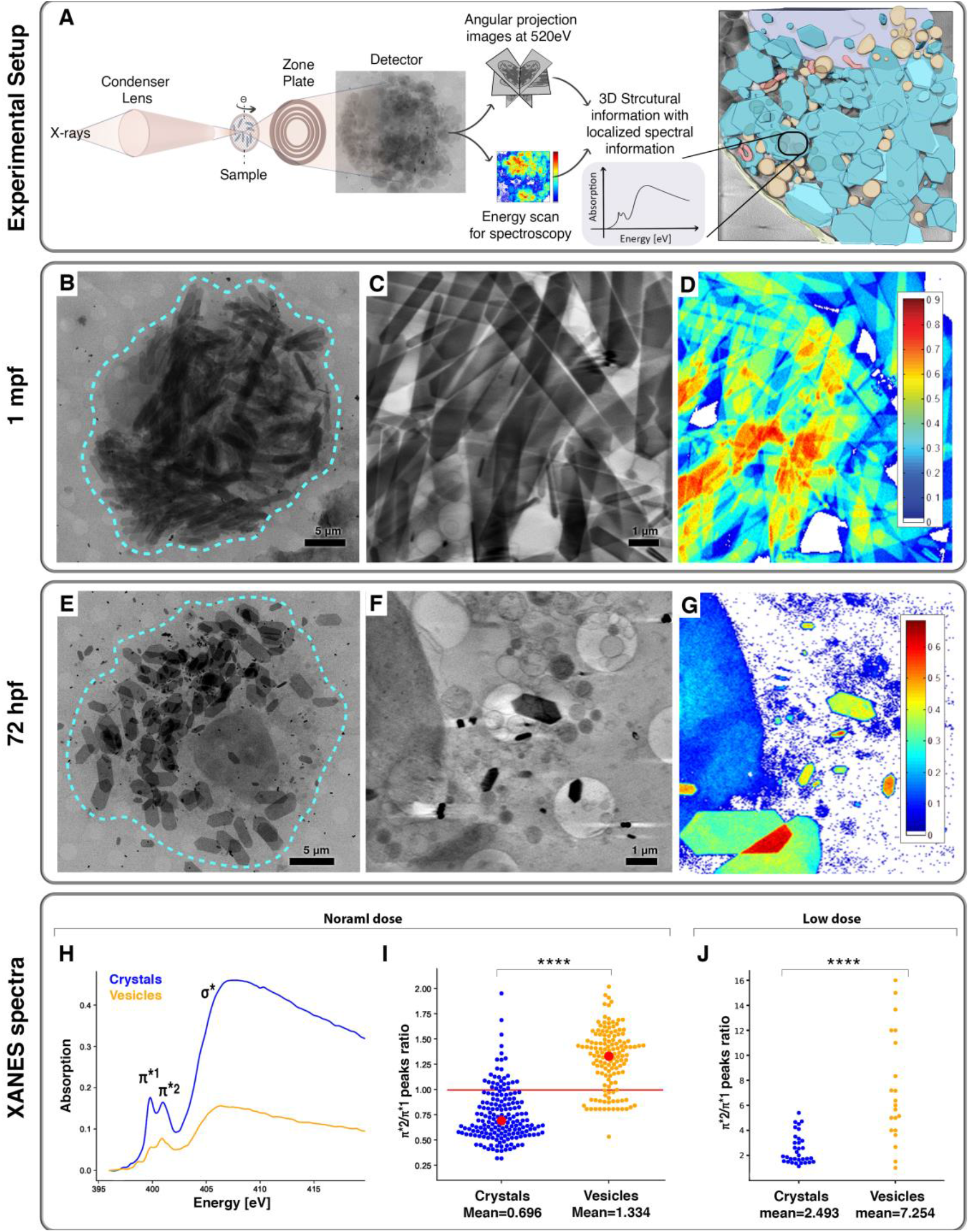
Cryo soft X-ray microscopy and XANES reveal amorphous guanine within intracellular vesicles. **(A)** Schematic illustration of the experimental design and data collection process using cryo-SXM, which enables correlated cryo-tomography and XANES. **(B, E)** Cryo soft X-ray microscopy images of isolated zebrafish larvae iridophores at 1 mpf (B) and 72 hpf (E). **(C, F)** Two-dimensional volume slices from a reconstructed cryo-SXT of a 1 mpf cell (C) and 72 hpf cell (F), showing multiple crystals and vesicles. **(D, G)** XANES maps showing the distribution of guanine within a 1 mpf cell (D) and 72 hpf cell (G). **(H)** A cryo-XANES spectra of the nitrogen K edge of a crystal (blue) and a vesicle (orange). **(I)** Swarm graph showing the ratio between π*^1^ to π*^2^ peaks of crystals (blue) and vesicles (orange), using normal radiation dose. T-test result = −17.8926, p-value = 1.454e-50. Mean ratio for crystals 0.7 (± 0.318, n= 191), whereas for the vesicles is 1.3 (± 0.325, n= 142). **(J)** Swarm graph showing the ratio between π*^1^ to π*^2^ peaks of crystals (blue) and vesicles (orange), using low radiation dose. T-test result = −5.924, p-value = 2.537e-07.

We proceeded to gather two-dimensional (2D) projection images of iridophores within the nitrogen K-edge energy range of 395-420 eV, where nitrogen absorption is at its peak (**Movie S4**). Following data acquisition, the XANES images were refined through flat field normalization and filtering processes (see materials and methods). This treatment resulted in detailed 2D ‘Nitrogen maps’ that highlight areas of concentrated nitrogen. In the analysis of cells from 1 month post fertilization (mpf) larvae, where crystallization is predominantly complete, we primarily observed mature, elongated crystals (**Fig. 3C-D**). In contrast, cells from 72 hpf larvae revealed smaller crystals in close proximity to nitrogen-rich organelles (**Fig. 3F-G**). The abundance of N-rich vesicles in 72 hpf samples was key in uncovering the chemical identity and characteristics of nitrogen in these vesicles. We derived this information from the XANES spectra corresponding to various regions of interest within the nitrogen maps. Notably, the spectrum associated with guanine crystals displayed two distinct peaks at 399 eV (π*^1^) and 401 eV (π*^2^). These peaks represent the excitation of guanine nitrogen’s 1s electrons to π* orbitals. Additionally, a broader peak around 407 eV was observed, indicative of the 1s to σ* orbital transitions on the different nitrogen sites in the guanine molecule.

Notably, the darkly contrasted intracellular vesicles also displayed an X-ray absorption pattern akin to that of guanine, suggesting guanine as their primary component (**Fig. 3H**). However, a more detailed analysis of the vesicles’ absorption spectra revealed distinct variances when compared to the spectra of guanine crystals (**Fig. 3I-J**). Specifically, the π*^1^ and π*^2^ absorption peaks in the vesicles were broader than in crystals (**Fig. S3**), and their relative intensities differed significantly. This suggests a less ordered molecular arrangement of guanine within these vesicles, distinct from its ordered arrangement in the crystals (**Fig. 3H**). By overlaying the spectra of several hundred crystals and contrasting them with a similar number of vesicle spectra, a clear discrepancy in the ratios of the π*^1^ and π*^2^ peaks was observed. After normalizing these spectra, we calculated the ratio of the π*^1^ and π*^2^ peaks and found that, on average, this ratio for the crystals was 0.7 (± 0.318, n=191), whereas for the vesicles, it was significantly higher, at 1.3 (± 0.325, n=142, p=1.45*10^−50^) (**Fig. 3I**).

Building on the finding that guanine aggregates in the vesicles appear less ordered, we hypothesized that the X-ray absorption of the highly anisotropic guanine crystals would be markedly influenced by their orientation relative to the beam, while the absorption in vesicles would be comparatively unaffected. To investigate this, we conducted absorption measurements on both vesicles and crystals during sample tilting. Our results confirmed that the absorption by crystals was significantly dependent on their alignment with the beam (**Fig. S4**). This effect was especially pronounced when the planar crystals were oriented perpendicular to the beam, when we observed a substantial reduction in their π* peaks in parallel to an increase in their σ* peak. In contrast, the vesicles’ absorption remained relatively stable regardless of the tilting angle (**Fig. S5C-D**). These findings further support the idea that the guanine molecules in the vesicles are in a less ordered state.

In the course of conducting tilt series experiments, we noticed a notable change: extended radiation exposure seemed to modify the absorption spectrum of the crystals. Intrigued by this observation, we embarked on a more detailed investigation into the impact of radiation damage. We exposed both crystals and vesicles to repeated radiation and carefully observed the subsequent alterations in their absorption properties. For vesicles, the impact was straightforward - the amplitude of both π* and σ* absorption peaks diminished with increased radiation exposure (**Fig. S5B**). However, the response in crystals was distinctly different. While the σ* peak’s amplitude decreased due to radiation, the amplitudes of the π*^1^and π*^2^ peaks in the crystals surprisingly increased (**Fig. S4F and Fig. S5A**). We speculate that this increase in π* absorption could be due to radiation damage that induces bond-breaking and an overall reduction in electron density therefore altering the transition probability into anti-bonding π states in the XANES signal of nitrogen. This may change the XANES fine structure, it is also possible that the breaking of bonds between the molecular sheets allows molecules to rotate more freely thus affecting their orientation to the X-ray beam.

To investigate if the differences in the π*^2^ to π*^1^ peak ratios in crystals compared to vesicles we observed were only a result of differential responses to beam damage, we conducted a series of experiments designed to minimize radiation effects, making sure that no visible damage is apparent in the course of a single scan (materials and methods and **Fig. S6**). We measured both crystals and vesicles using a low-dose radiation approach. Under these conditions, the difference in the π*^2^ to π*^1^ ratios between crystals and vesicles was less pronounced (**Fig. 3J**). However, both the π*^2^ to π*^1^ ratios and their spread were significantly larger in the vesicles. Considering that the vesicles orientations do not affect their peak ratios, these observations suggest that factors beyond mere molecular ordering influence the π*^2^ to π*^1^ ratio. Previous studies on the XANES characteristics of guanine indicated that different protonation states of guanine show different absorption profiles^43^. Specifically, it was shown that protonated guanine exhibits an elevated π*^2^ to π*^1^ ratio compared to the neutral form. To further test this hypothesis, and to compare aspects of the expected spectra of neutral and protonated forms of guanine, we conducted a series of DFT calculations for the density of empty states. The results, shown in **Fig. S7**, compare the calculated density of states (DOS) of (a) a gas phase guanine molecule, and (c) a gas phase N-7 protonated guanine molecule (we restrict our attention to the DOS, as opposed to a full simulation of XANES data, in order to avoid controversies arising from different levels of theoretical treatment of core holes). Considering the ratio between the first two peaks as a proxy for the π*^2^ to π*^1^ ratio, we find that it is ∼1.2 for the isolated guanine molecule, but ∼1.75 for the protonated form. This supports the experimental findings on the difference between the ratio of π*^1^ and π*^2^. Thus, the differences in the π*^2^ to π*^1^ ratio observed for the vesicles could indeed suggest that the vesicle microenvironment is also acidic, and that during organelle maturation and crystal formation, the pH of the organelle gradually neutralizes.

To test if the iridosomes do provide an acidic environment, we performed confocal imaging with LysoTracker and LysoSensor dyes^44,45^ known to selectively stain acidic intracellular compartments like lysosomes. Our observations during the early developmental stages (48-96 hpf), a period characterized by active crystal formation before the crystals reach full maturity, revealed strong LysoTracker staining of iridosomes (**Fig. 4A and Fig. S8A**), corroborating the acidity of these organelles. In contrary, at later developmental stages (7 days post-fertilization), the iridosomes showed markedly weaker staining (**Fig. S6A**). Importantly, within the same cell at early developmental stages, iridosomes at varying levels of maturation exhibited differential LysoSensor staining, with smaller, less mature iridosomes displaying more pronounced staining (**Fig. 4B and Fig. S8B**). We then systematically quantified the LysoSensor staining of crystals at different developmental stages in different larvae, and found a similar trend, smaller iridosomes showed stronger signal (**Fig. 4B**). These findings collectively reinforce our earlier results, suggesting that iridosomes are initially acidic and that the organelle’s pH gradually neutralizes as it matures, and crystal formation ensues.

**Figure 4:**
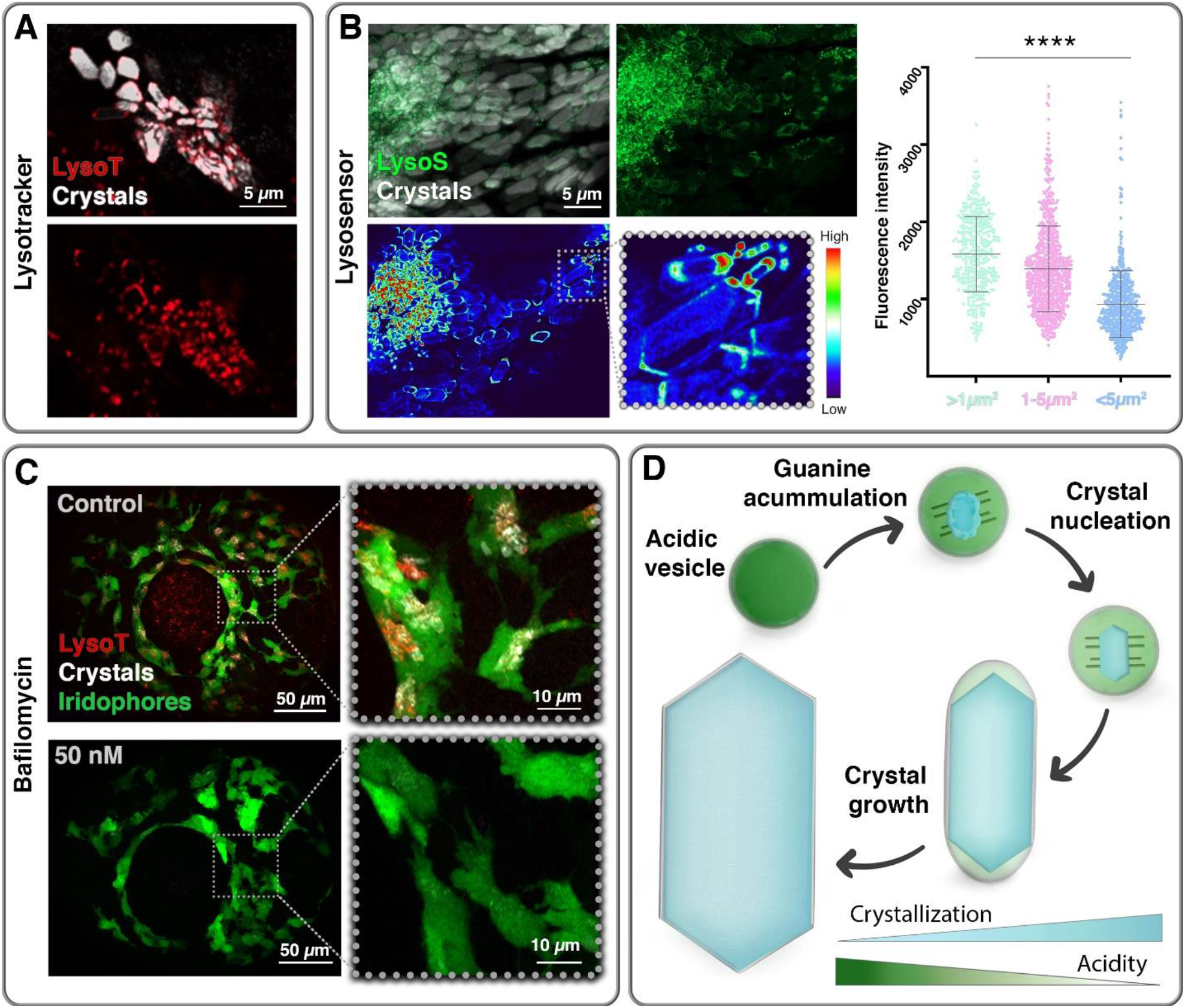
pH alterations drive guanine accumulation and crystal formation within the iridosome. **(A)** Confocal microscopy images of zebrafish larvae at 48 hpf stained with LysoTracker showing iridophore intracellular acidic compartments (red) and crystals (white) within the iridophores. **(B)** Confocal microscopy images of zebrafish larvae at 96 hpf stained with LysoSensor, which exhibits increased fluorescence in acidic environments. Top – maximum intensity projection image from a Z-stack showing iridophore intracellular acidic compartments (green) and crystals (white) within the iridophore. Bottom - heatmap representation of a single plane of the Z-stack from the top panel. Right–swarm plot showing the fluorescent intensity of small, under 1 μm^2^ iridosomes (green) compared to medium 1-5 μm^2^ (pink) and large over 5 µm^2^ iridosomes (blue). Mean ± SD is shown. ****p < 0.0001, two-tailed, unpaired student t-test. (**C)** Confocal microscopy images of 60 hpf zebrafish larva iridophores (green, transgenic TDL358:GFP) treated with DMSO only (top), or treated with 50 nM Bafilomycin A1 (bottom). Lysotracker staining (red), crystals (white). **(D)** A proposed model for guanine accumulation and crystal formation. Initially, the iridosome is highly acidic, allowing soluble guanine to accumulate in a protonated form. As the iridosome matures, its pH gradually transitions toward neutrality, prompting the deprotonation of guanine, which triggers crystal nucleation. The continued gradual increase in pH promotes crystal growth by shifting the equilibrium toward neutral guanine, providing additional building blocks for the growing crystals.

To further assess the importance of acidity in guanine accumulation and crystal formation, we administered Bafilomycin A1, an inhibitor of the vacuolar-type ATPase (V-ATPase)^44^, to larvae at various developmental stages (ranging from 36-54 hpf). Larvae subjected to this treatment exhibited a marked reduction in LysoTracker staining in iridophores without affecting the overall development (**Fig. S8C**). Moreover, these larvae displayed a significant reduction in crystal quantity compared to untreated controls (**Fig. 4C**). These findings imply that the acidic environment within iridosomes is crucial for the accumulation of guanine and the subsequent formation of guanine crystals.

## Discussion

The precision of guanine crystal formation within cells requires exact control over the solubility of guanine. In our study, we demonstrated that cells accomplish this through the careful regulation of the microenvironment within the iridosome, the organelle in which crystals form. At first, these organelles are highly acidic, which allows soluble guanine to accumulate in a protonated form. As iridosomes mature, they transition toward a more neutral pH, prompting the deprotonation of guanine, which in turn triggers crystallization. Additional gradual increase in pH promotes crystal growth by further shifting the equilibrium toward neutral guanine, supplying the growing crystals with additional building blocks (**Fig. 4D**).

We first showed that zebrafish larval iridophores are populated with numerous nitrogen-rich vesicles, measuring between 100 to 500 nm, alongside small guanine crystals. Cryo-XANES analysis correlated with cryo-SXT 3D imaging revealed that the nitrogen-rich material within these vesicles is guanine. However, the X-ray absorption characteristics of the vesicles, namely, their markedly low dependence on orientation and their response to beam damage, suggest that the guanine within them lacks the ordered structure observed in the crystals. This was further substantiated by the absence of electron diffraction patterns from the vesicles, affirming the non-crystalline nature of the encapsulated guanine.

Guanine is particularly insoluble in aqueous solutions, and under ambient conditions there is a strong driving force towards crystallization. Thus, the presence of concentrated amorphous guanine within vesicles could only be obtained under conditions that increase its solubility and inhibit its crystallization^18^. The mechanisms that enable the trafficking and accumulation of insoluble guanine within the intracellular vesicles and the driving force behind crystallization have yet to be illuminated. By applying a multidisciplinary approach integrating state-of-the-art electron microscopy, spectroscopy, live imaging, and pharmacological perturbations we show that the vesicle microenvironment is acidic and that during organelle maturation and crystal formation, the pH of the organelle gradually neutralizes. This implies that pH changes enable both guanine accumulation and drive crystallization within the iridosomes.

Firstly, the elevated π*^2^ to π*^1^ ratios we observed for the vesicles compared to the crystals suggest that the vesicles are acidic. The spread of π*^2^ to π*^1^ ratios exhibited by different vesicles even when using a low-dose radiation approach suggests that organelles at different maturation stages have different pH values. Using live imaging, we further demonstrated, that early iridosomes are highly acidic, and that during maturation they gradually become less acidic, and subsequently driving the growth of the guanine crystals within them.

It was previously suggested that iridosomes like other pigment organelles such as melanosomes are lysosome-related organelles^16,21^. One of the characteristics of LROs is their low pH. Our findings that the iridosomes are acidic as well, further support this classification. Recent studies have illustrated that guanine crystals develop through the templated nucleation of thin leaflets on preassembled scaffolds^31,32^. Studies on other functional amyloids indicated that pH fluctuations can reversibly influence amyloid assembly^46^. Thus, the initially acidic environment within the iridosome may facilitate the formation of amyloid fibers, which could template the heterogeneous nucleation of the guanine crystals. The subsequent neutralization of pH as the organelle matures and the crystals grow, may lead to the disassembly of these amyloid structures, offering potential reason for their absence in developed organelles.

Finally, we found that inhibiting vacuolar V-ATPase, via Bafilomycin A, reduced the acidity of the iridosomes and led to a noticeable decrease in crystal formation. Prior investigations into the iridophores’ proteome and transcriptome^47^ revealed the upregulation of several V-ATPase genes, including tcirg1a and tmem179b, suggesting the involvement of multiple proton pumps in regulating iridosome acidity. This indicates a complex mechanism behind the precise adjustment of the iridosome’s microenvironment. Future research will be essential to pinpoint the specific proton pumps responsible for this regulation in iridophores.

In conclusion, our findings provide new insights into the molecular mechanisms that enable guanine accumulation and drive crystallization within iridosomes, highlighting the critical role of pH gradients in the precise orchestration of biogenic crystal formation.

## Acknowledgments

This work was supported by an ERC Starting Grant (Grant number: 101077470, “CRYSTALCELL”) and by the Israel Science Foundation (grant No. 691/22) awarded to D.G. This work benefited from access to ALBA and has been supported by iNEXT-Discovery, project number 871037, funded by the Horizon 2020 program of the European Commission. Cryo soft X-ray experiments were performed at the MISTRAL Beamline at ALBA Synchrotron with the collaboration of ALBA staff. We thank Dr. Eva Pereiro for her support and assistance in coordinating with the synchrotron. Additionally, we thank Dr. Tali Lerer-Goldshtein for her support in developing the project’s experimental approach. Electron microscopy studies were conducted at the Irving and Cherna Moskowitz Center for Nano and Bio-Nano Imaging at the Weizmann Institute of Science.

